# Autofocusing molecular simulations for integrative structural biology of broad conformational ensembles

**DOI:** 10.64898/2026.10.09.757932

**Authors:** Hanne S. Antila, Markus S. Miettinen

**Affiliations:** Department of Theory and Bio-Systems, Max Planck Institute of Colloids and Interfaces, 14424 Potsdam, Germany; Department of Biomaterials, Max Planck Institute of Colloids and Interfaces, 14424 Potsdam, Germany; Department of Physics, Freie Universität Berlin, 14424 Berlin, Germany; Department of Chemistry, University of Bergen, 5007 Bergen, Norway; Computational Biology Unit, Department of Informatics, University of Bergen, 5008 Bergen, Norway

**Author notes:** **Correspondance** Correspondence and requests for materials should be addressed to M. S. M.

## Abstract

Integrative structural biology seeks molecular descriptions consistent with complementary experimental measurements. A particular challenge arises for molecules whose broad conformational ensembles resist description by a few representative structures, such as natively unfolded proteins and fluid-phase biomembrane lipids. Although molecular simulations can generate such ensembles, their simultaneous agreement with a large number of experimental constraints is commonly prevented by model inaccuracies. Here we introduce “autofocusing” as an approach that integrates experimental information by automated finetuning of the physical model generating the ensemble. We demonstrate it for a fluid bilayer of the phospholipid POPC, for which no existing force field has simultaneously reproduced the complete set of 82 NMR C–H bond order parameters. Autofocusing brings all the 82 within experimental uncertainty while preserving agreement with independent small-angle X-ray scattering measurements and retaining or improving agreement with untargeted NMR relaxation rates and effective correlation times, thus supporting its description of membrane structure and dynamics beyond the calibration targets. The resulting simulations challenge a long-standing two-state interpretation of phosphatidylcholine headgroup structure, instead supporting a broad conformational ensemble poorly represented by crystal-derived conformers. These results establish autofocusing as a practical approach to constructing experimentally grounded conformational ensembles while retaining the computational tractability of classical molecular dynamics.

## INTRODUCTION

The classic paradigm of structural biology takes functional information to be encoded in well-defined conformations of folded biomolecules. It is becoming increasingly clear, however, that key biomolecules of cellular function—such as proteins [1, 2], RNA [3], lipids [4–6] and polysaccharides [7]—often lack a single stable conformation under physiological conditions. Intriguingly, this structural promiscuity does not mean that the exact population of conformations (and the exact inter-conformation dynamics) would not be crucial for such molecules’ correct biological function: These structural chameleons often play roles in regulation [8, 9], recognition [10, 11], and signalling [12, 13], where minute differences in atomic level structural details can have extreme consequences [14–16]. The structural variability appears, in fact, to be high in information content.

For stably folded biomolecules, we have grown accustomed with experimental visualisations at atomic resolution [17–25], but for the populations of intrinsically unstructured biomolecular conformations—and the conformational fluxes within these populations—a similar sub-nano(meter and second) accuracy is lacking. Consequently, the unstructured half of the biomolecular universe remains largely invisible, and our ability to investigate how information is coded in the structural variability severely limited.

Since 1970s, it has been visioned that the sub-nano realm could be seen by using physics-based simulations as a “computational microscope” [26–35]. But, as the direct use of quantum mechanics continues to be prohibitively expensive for biomacromolecular spatiotemporal scales (*>* 1 000 atoms over *>* 10 ns), and its classical approximation molecular dynamics (MD) simulations has not been able to faithfully reproduce atomistically sensitive data (e.g., from nuclear magnetic resonance (NMR) spectroscopy [36–39] and scattering [40–42]), it remains unclear if true computational microscopy can, in practise, be achieved.

Realizing the vision of sub-nano-level computational microscopy through MD is posed with substantial challenges and open questions. First, can any classical approximation retain sufficient realism to veritably describe an intrinsically disordered biomacromolecular system? Second, tuning hundreds of heavily cross-correlated non-linear parameters to match dozens of experimental data points simultaneously is humanly daunting, if not impossible. Can it be automated? By successfully performing automatic data-driven calibration of an existing MD force field, we tackle both of these questions. Our demonstration achieves veritable computational microscopy of an intrinsically disordered biomolecular system—a milestone on the path to universal computational microscopy for structural biology.

In reaching this milestone, we show that computational microscopy is achievable for a real-world intrinsically disordered biomolecular system and is not precluded by the classical approximation. As the computing cost of our autofocusing remains feasible, we also demonstrate how atomistically-sensitive but hard-to-interpret experimental (NMR) data can now be given an intuitive subnano-resolution “microscopic” interpretation. The benchmark we provide has an immediate application in revealing the structure and dynamics of a model biomembrane system. Other initial applications are expected to arise in integrative structural biology of intrinsically disordered biomolecules: An MD force field calibrated to simultaneously match a spectrum of atomistically-sensitive experiments (NMR spectroscopy, X-ray scattering, and cryoelectron microscopy, for example) will provide a computational microscope that unifies all these methods. However, using MD to realize the full promise of computational microscopy—the realistic out-of-the-box simulation of any biomolecular system—will still require a force field that is not system-specific but transferable.

Our demonstration of autofocusing for integrative structural biology also paves the way towards a universal transferable force field, as for the successful simultaneous matching of large sets of data for different molecules and various external conditions, automated calibration will be paramount.

### A SUITABLE BIOMOLECULE

To demonstrate veritable Ångström-resolution computational microscopy, we perform automated calibration of MD simulations for a non-trivial intrinsically unstructured biomolecular system: a bilayer of POPC (1-palmitoyl-2-oleoyl-sn-glycero-3-phosphocholine) phospholipids (see Fig. 1B). Lipid bilayer biomembranes are particularly suitable proof-of-principle systems as (i) in their biologically relevant (liquid disordered) state lipids are intrinsically unstructured; (ii) a single lipid is reasonably small (~ 10^2^ atoms), and thus amenable for automated calibration; and (iii) a bilayer simulation must contain at least a few dozen lipids, thus naturally providing parallel sampling. Most important, however, is (iv) the existence of high-resolution structurally-sensitive experimental data that can be calculated from MD simulations directly [45]; these are the NMR-measured C–H bond order parameters:

**FIG. 1.**
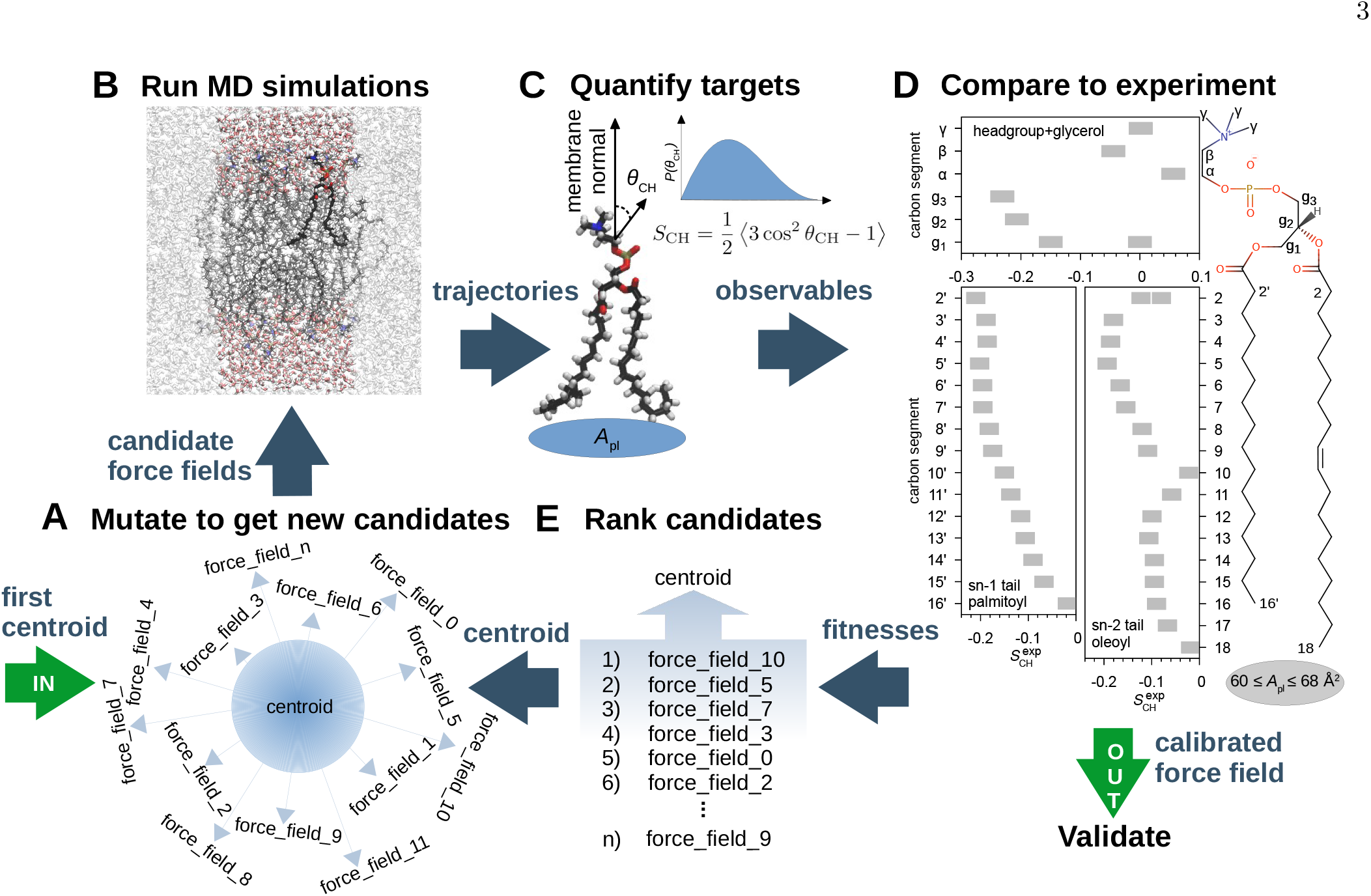
Autofocused Å-resolution computational microscopy of intrinsically disordered biomolecular matter. A) We use an evolutionary algorithm [43] to create *n* new MD force field candidates by mutating the parameters of a centroid individual. The first centroid is the progenitor force field, here Slipids [44]. B) With each candidate, we conduct an MD simulation; here on a 34-lipid bilayer patch (in periodic boundary conditions: the system shown in colour, its periodic images in gray). C) From the simulation trajectories, we extract the target observables; here the area per lipid *A*_pl_ and the 82 C–H bond order parameters *S*_CH_ (see Eq. 1). D) We compare against the experimental ranges of target observables (grey) to obtain (see Eqs. 2 and 3 in Methods) the candidate fitnesses. E) We rank the *n* candidates according to their fitnesses, and use the centroid of the ^*n*^/_2_ fittest to create the next generation of candidates.

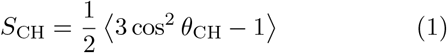

where *θ*_CH_ is the angle between the C–H bond of interest and the biomembrane normal, and the average is over the simulated lipids and time (see Fig. 1C). A POPC lipid has 82 C–H bonds (see Fig. 1D), giving 82 unique *S*_CH_— an excellent target set [38, 46] for automated calibration. In addition to these target data, a rich set of independent high-resolution validation data exist for POPC; for example small angle (x-ray and neutron) scattering data are sensitive for membrane dimensions such as thickness and area per lipid [47], and NMR relaxation data for individual lipid dynamics [48, 49].

Our goal is to achieve a force field that reproduces within experimental accuracy all the 82 *S*_CH_ in POPC. This is a difficult task because the mapping of an essentially quantum mechanical system to a classical one will necessarily be imperfect, and the classical force field parameters are highly sensitive, cross-correlated, and have nonlinear effects on the biomolecular structure. To be able to perform computational microscopy of biomolecular systems at will, we need an automated approach to tune the force field parameters such that a high-fidelity match with the experimental target data is reached.

### THE HIGH-FIDELITY FORCE FIELD CALIBRATOR

We designed an automated calibrator that iterates MD force field parameters until the complete set of experimentally-known constraints is met (Fig. 1). For structural biology, the key advance of such autofocusing is achieving automatic integration of a vast amount of experimental knowledge into a single, consistent, physics-based spatiotemporal model. The set of calibration targets may contain any experimental observables calculable from MD, including but not limited to: NMR relaxation rates [45], scattering intensities (also from x-ray free electron laser [50–52] experiments), and electron microscopy electron density maps [53, 54].

Here we use 83 different experimentally known targets: the 82 C–H bond order parameters *S*_CH_ (see Eq. (1)) and the area per lipid *A*_pl_. We require the *S*_CH_ to fit within their experimental uncertainty (*±* 0.02) [38] and the *A*_pl_ within the spread of values (60–68 Å^2^) reported across various indirect experiments [47, 55, 56]. The resulting high-dimensional nonlinear calibration problem we tackle with the Covariance Matrix Adaptation Evolution Strategy (CMA-ES) [43]. We discuss the key points below; see also Methods.

Autofocusing is initiated by inputting an existing force field as the first ‘centroid’ (Fig. 1: ‘IN’); we performed calibrations starting from the Slipids [44] force field known to be among the best existing models in capturing the true conformational ensemble (*S*_CH_) [38, 57] and dynamics [49] of our intrinsically unstructured proof-of-principle system. From the centroid, a number of candidate force fields are created by mutating (i.e., random perturbation of) the force field parameters (Fig. 1A). With each of the resulting force fields, an MD simulation (Fig. 1B) is run, and the target observables calculated (Fig. 1C). Comparison against the target experimental values (Fig. 1D) gives the fitness of each candidate, based on which they are ranked (Fig. 1E). The centroid of the fittest candidates is then used to create the next generation of candidate force fields (Fig. 1A).

When a candidate reproduces the complete set of experimental targets (Fig. 1: ‘OUT’), it is validated against non-target data. This crucial final step in autofocusing a high-fidelity force field alleviates the risk of overfitting to the chosen targets, and ensures that experimental observables outside the targets are also well represented. Here we used the form factors *F* (*q*) from small-angle X-ray scattering (SAXS), as well as the spin-lattice relaxation rates *R*_1_ and the effective correlation times *τ*_e_ from ^13^C-NMR, to validate that both the overall bilayer structure and the intramolecular dynamics remain well described in the autofocused force field.

### THE AUTOFOCUSED FORCE FIELD

Figure 2 shows the outcome of a force field calibration. Observables calculated from 1-*µ*s MD simulations using the successful candidate (598-3) and the state-of-the-art force field used to initiate the autofocusing (Slipids [44]) are compared against NMR and SAXS experiments. The autofocused model faithfully reproduces all the 82 C– H bond order parameters *S*_CH_ (see Eq. (1)) within experimental accuracy (Fig. 2A); this gives these hard-to-interpret atomistically-sensitive data an intuitive visual exposition, effectively manifesting Å-resolution computational microscopy.

**FIG. 2.**
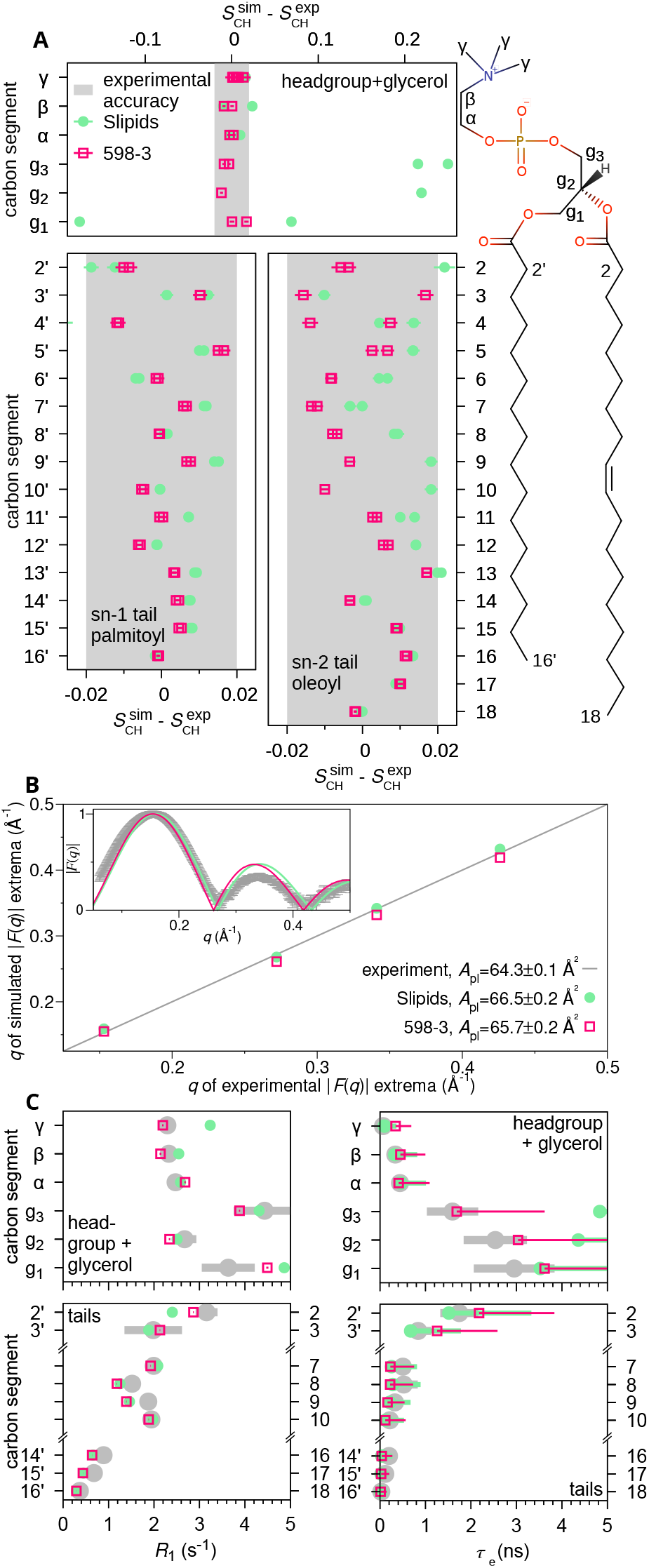
Autofocusing robustly captures Å-scale detail. Calibrated simulation assessed against the targets (A), and validated for overall bilayer structure (B) and conformational dynamics (C). A) In contrast to the progenitor model (Slipids [44], filled circles), all the 82 *S*_CH_ (see Eq. 1) of the autofocused model (598-3, open squares) match the experimental *S*_CH_ (from Refs. 38, 46) within experimental uncertainty (gray shaded). B) Experimental (from Ref. 47) vs. simulated extrema locations of SAXS form factor curves |*F* (*q*) | along with the areas per lipid *A*_pl_ (legend); on the dashed line *x*=*y*. Inset shows the full |*F* (*q*) | scaled by the first lobe height. C) Validation against experimental (grey, from Ref. 58) ^13^C-NMR spin-lattice relaxation rates (*R*_1_, left) and effective correlation times (*τ*_e_, right) show improvement compared to the progenitor model both at the~ 1-ns time scale (*R*_1_) and across time scales (*τ*_e_); note that both tails contribute to the data for segments 2’–3’ (2–3 in the sn-2 chain) and 14’–16’ (16–18).

Figure 2B validates that the calibration preserved well the overall bilayer structure. Shown is the comparison of the zero-point and extrema locations in the simulated and experimental SAXS form factor curves *F* (*q*): The data remain well reproduced by the autofocused model. The inset shows the full *F* (*q*), i.e. the Fourier transform of the electron density profile along the bilayer normal as obtained from scattering experiments: unsigned and on a relative scale [45].

Figure 2C validates the conformational dynamics. Shown are comparisons to the experimental ^13^C-NMR spin-lattice relaxation rates *R*_1_ (left) and effective correlation times *τ*_e_ (right) [48, 49]. The *R*_1_ are sensitive to dynamical processes of the C–H bond orientation in a narrow NMR-setup-dependent window (here ~ 1 ns), whereas the *τ*_e_ are composite observables that capture C–H bond dynamics across a range of 10^*™*13^– 10^*™*6^ s. Agreement is good both when focusing on the nanosecond window (*R*_1_) and when considering the whole range of processes (*τ*_*e*_). In fact, even though dynamics was not a calibration target, the calibrated model outperforms the state-of-the-art force field (see, e.g., *τ*_e_ for the g_3_ and g_2_ carbons): The autofocused MD simulation not only samples the correct conformational ensemble, but does so with almost exactly correct dynamics.

As summarised by Fig. 2, veritable Å-resolution computational microscopy is achievable for a nontrivial intrinsically disordered biomolecular system by autofocusing classical MD force fields—and this is not conditional to overfitting that would destroy the conformational dynamics or the overall properties of the system. We proceed to see what computational microscopy reveals on biomembrane structure and dynamics.

### BIOMOLECULAR INSIGHT

Biomembranes are remarkable functional materials that despite their simple core design can host rich functional complexity. They can be highly permeable to select substances while being practically impenetrable to most; they are also at the same time mechanically stable as well as fluid and flexible. These various seemingly contradictory abilities enable vital cellular functions such as vesicle budding and fusion, the spatial organization of biochemical reactions, and signal transduction—all of which have their origins in the structural and dynamic details of the biomembrane’s lipid constituents. Consequently, accurate molecular-level knowledge on membrane lipids provides mechanistic insights into numerous cellular processes. Due to the liquid-disordered state of the biologically relevant phospholipid bilayers, however, detailed conformational resolution has remained unattainable, even with advanced microscopy, scattering, and spectroscopic techniques. Here, we demonstrate the use of the autofocused computational microscope by resolving, with Å-level fidelity, the conformational dynamics of POPC, a key phospholipid in eukaryotic cell membranes.

We begin with the PC headgroup. The current understanding (see Refs. 60–62 for review of phospholipid headgroup structure and dynamics) remains heavily influenced by the number of conformers (two for PC, see

Fig. S1) occurring in X-ray crystal structures. These frozen conformations are utilized to extrapolate headgroup structural characteristics also in the biologically-relevant liquid-disordered state. Specifically, NMR C–H bond order parameter (*S*_CH_, see Eq. (1)) data in the liquid-disordered state have been interpreted through this template, i.e., to suggest a rapid exchange between the two principal crystalline conformers. Our MD simulations autofocused to reproduce these same NMR data, however, provide little support for this two-state interpretation. While the two principal crystalline conformations do occur, the conformational space of PC headgroup is not sufficiently described by them: Within *±* 30^*°*^ windows of those two lie less than 5% of all the PC head-group conformations (see Table S1 for dihedral-spesific data). Instead, the true conformational ensemble of PC headgroups in the biologically-relevant liquid-crystalline state is seen to comprise a broad continuum of conformations, all of which are easily accessible with thermal energy (Fig. 3A). As a specific example, Fig. 3D–F show a path joining the two principal crystalline conformers. Notably, a path with similarly low barrier could be drawn between any states in the free energy map (Fig. 3A), underscoring that there are no regions that would be separated by much-higher-than-thermal barriers.

**FIG. 3.**
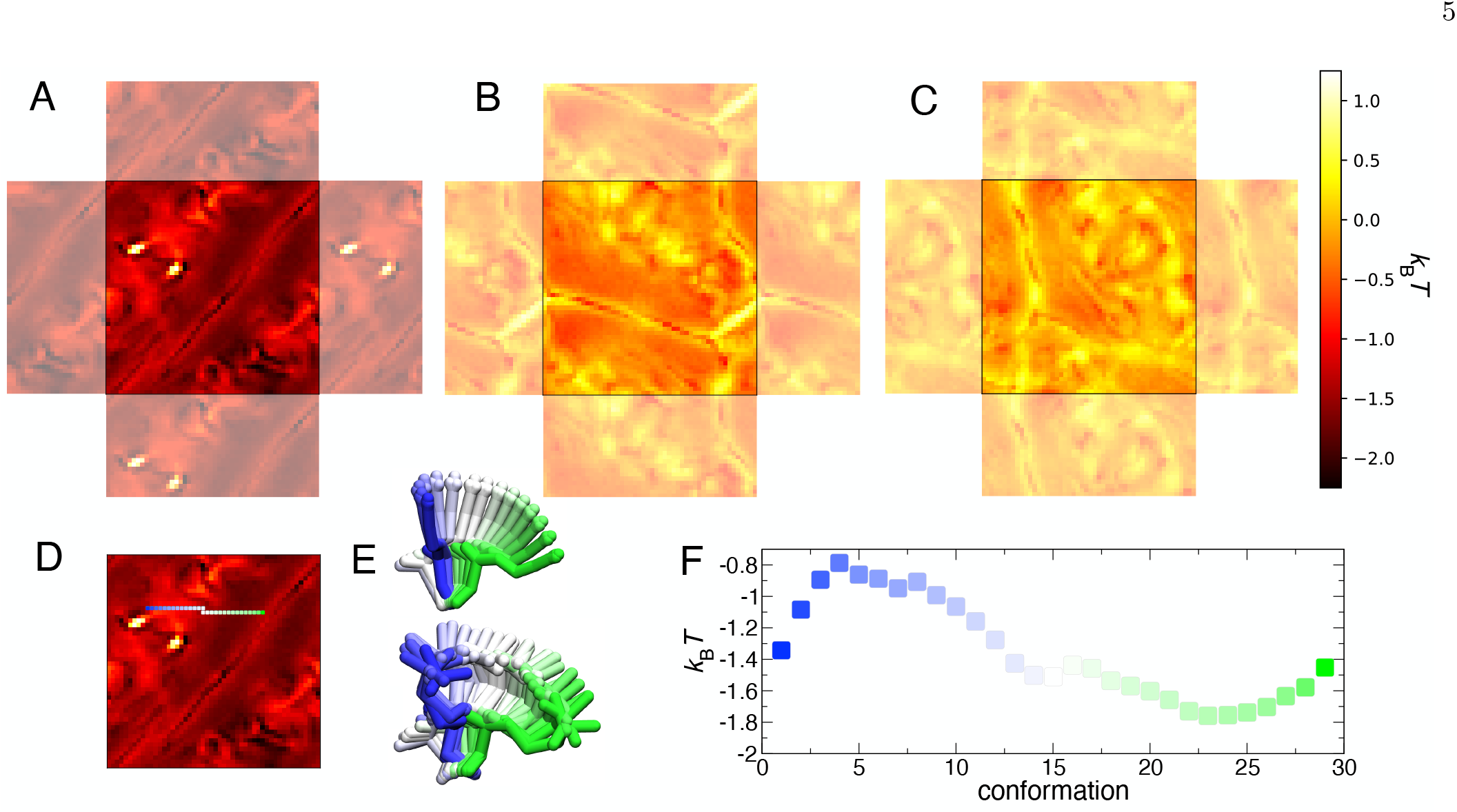
Autofocused simulation reveals PC-headgroup conformations as a thermally accessible continuum, not discrete states. The conformational free energy space of POPC visualized as self-organized maps (SOMs). The molecule is divided into three fragments: A) headgroup+glycerol, B) sn-1 tail, and C) sn-2 tail; the SOMs have periodic boundary conditions visualized with shaded areas. D) A 29-step conformation path drawn on the headgroup SOM from one crystal [59] conformation (step 1, blue) to another (step 29, green). E) The average SOM cell conformations (top) and the corresponding best-matching conformations from the MD trajectory (bottom) colored according to the path step. F) The free energies along the drawn path.

For the glycerol backbone, the autofocused simulations again highlight the difference between the lone frozen crystal structures and the wide continuum conformational ensemble of the liquid-disordered state (Figs. S3, S2). Focusing on the orientational distribution of the bond vectors starting from the central backbone carbon C^g2^, we observe that while the C^g2^–C^g3^ bond does, as in the crystalline conformers, tend to orient nearly parallel with the membrane normal (Fig. S3A), for C^g2^–C^g1^ and C^g2^–O^g2^ there is a qualitative switch of roles: In the liquid-disordered state it is more often the C–C bond that lies in the membrane plane, while C–O is the one inclined slightly more towards the membrane interior (Fig. S2A).

The latter difference challenges the most apparent shared feature in the few successfully solved phospholipid crystal structures: While the *sn*-1 fatty acid chain, “the leading tail” [60], dives directly into the membrane from its attachment to the glycerol backbone C^g2^, the other bends at a 90-degree angle (Fig. S1). As indicated by Fig. S2A, the “leadership” of the *sn*-1 tail that is seen in the crystal structures is much less conspicuous in the liquid-disordered state. Also the *sn*-2 tail is typically not bent 90^*°*^, but the O^g2^–C1 bond inclines towards the membrane interior (Fig. S5E) almost as much as the O^g1^–C1’ bond does (Fig. S5D). Nevertheless, also in the autofocused simulations the *sn*-1 more often assumes the leading role (Fig. S5C,F). As for the headgroup, also in the tail SOMs (Figs. 3B,C and S9B,C) one can traverse from any conformer to any other with the cost of thermal energy. For more detailed discussion comparing the customary interpretation of experiments in the literature and the conformers we observe in the autofocused simulations, see the supplementary material.

Let us end with a glimpse of the dynamics. Figure S8 shows the tail dihedrals in a single POPC molecule as a function of time over a 25-ns window. In agreement with the trend apparent in the effective correlation times (Fig. 2C), the isomerization rates increase towards the tail ends. The characteristic lifetimes of the gauche (indicated as red in Fig. S8) states are typically well below 100 ps, with the exception of the dihedrals neighbouring the double bond (*β*_8_ and *β*_12_), whose gauche states are sufficiently long-lived for visible breaks to emerge in the trans-occupation (black): Although the *cis* double bond does perturb the tails and thus create more space for conformational changes, locally it restricts dihedral isomerization. This is most clearly seen in the *β*_8_ dynamics, which are even slower than those of the *β*_7_. Towards the tail ends, the isomerization dynamics of the two tails then become increasingly similar, i.e., not affected by the unsaturation.

## CONCLUSION

Autofocused computational microscopy can generate broad conformational ensembles that simultaneously satisfy a large number of experimental constraints, thus providing a practical approach to integrative structural biology. As demonstrated for a fluid lipid membrane, automated finetuning of a tractable classical force field not only achieves simultaneous agreement with the structural targets but also preserves or improves agreement with independent measurements of membrane structure and dynamics. Autofocused simulations can therefore be a valuable tool for exploring the natively disordered regions of structural biology, whose conformational heterogeneity challenges both structural characterization and intuitive interpretation. We exemplified this through reassessment of the conformational ensemble of biomembrane lipids: The autofocused simulations reproduce data conventionally interpreted as arising from rapid exchange between a few crystalline conformers, but in fact reveal a broad conformational ensemble of which those conformers account for only a small fraction.

For near future, the increasing computing power offers the prospect that acquisition of new experimental data would be routinely accompanied by rapid automated refinement of force fields. Such on-the-fly autofocusing could fundamentally reshape the role of molecular dynamics simulations in revealing the structure and dynamics of biomolecular systems.

## Acknowledgments

The authors thank Dr. Tiago M. Ferreira and Dr. O. H. Samuli Ollila for useful comments. H. S. A. gratefully acknowledges funding from the Instrumentarium Science foundation, Osk. Huttunen Foundation, and the Alexander von Humboldt foundation. MSM is supported by the Trond Mohn Foundation BFS2017TMT01.

## Supplementary information

Supplementary information is available for this work containing further analysis of the lipid conformations from the simulations against the experiments.

## Data and code availability

The force field and trajectory resulting from the automated calibration is available in Zenodo [63] in full, whereas the optimization algorithm is provided in a GitHub repository [64] that links to an additional Zenodo repository [65] with data and analysis scripts used for the figures.

## Author contributions

H. S. A and M. S. M conceptualized the work together.

H. S. A devised the algorithm, conducted the simulations and data analysis. Both authors interpreted the findings and contextualized the results. H. S. A drafted the first version of the manuscript. M. S. M conducted further data analysis and wrote the final version of the manuscript together with H. S. A.

## METHODS

### Optimization

We use an evolutionary algorithm to calibrate the force fields to the target data. In this class of algorithms, at each round, a number of solutions (here: force fields) are generated. They are then ranked based of how well they solve the problem (fitness) and the best ones are used to generate a new set of solutions via mutation, recombination, or both, with the goal that the good properties of the solution are passed on to the next generation and the algorithm eventually converges to an optimum.

More spesifically, we use the (*µ/µ*_W_,*λ*) covariance matrix adaptation evolution strategy (CMA-ES) algorithm where *λ* denotes the number of solutions (force fields) generated at each optimization round, and *µ/µ*_*w*_ indicates that a weighted mean of the best solutions is used as the centroid individual, from which the *λ* solutions of the next generation are generated by the (adaptive) mutation, where the mutation strength is informed by the previous generation(s). Here we use *λ* = (4 +floor(3 log(*N*)) solutions, where *N* is the number of parameters to be optimized and the floor function rounds the number down to the closest integer. Number of *µ* = *λ/*2 best solutions are selected to generate the centroid for the next optimization round with the weights chosen as 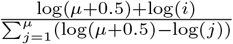, where *i* is the *I* individual when the solutions are ranked by their fitness and log denotes the natural logarithm. The initial standard deviation of the random distribution used for mutation is set at *σ*_ev_=0.15. More detailed description of the algorithm is available in Ref. [43]. The implementation of the algorithm was done based on the DEAP python library [66]. The parameters optimized for the Slipids-based model are the dihedral angle barrier heights: we tune all the dihedral barriers involving the atoms in the lipid head-group and glycerol backbone regions up to and including the C2 carbon (and the hydrogens attached to them) in the beginning of the lipid tails (139 parameters).

#### MD simulation within each optimization round

The fitness of each force field is calculated by extracting the area per lipid (*A*_pl_) and the 82 NMR order parameters (*S*_CH_) from an NPT molecular dynamics simulation consisting of 34 lipids (17 per leaflet) and 30 waters per lipid. The TIP3p [67] model is used for water. The simulation for each force field is first equilibrated for 3 ns using reduced timestep of 0.001 ps after which a production run of 35 ns, using timestep 0.002 ps, is conducted to extract the fitness. The initial configuration for simulating each force field is obtained from the preceeding optimization round, from the force field that was the closest (in terms of the 2-norm of the parameter values) to the current one being simulated. In the simulations, the temperature is kept at 300 K using the Nose– Hoover [68, 69] thermostat and the pressure at 1 bar with the Parrinello–Rahman [70] barostat and semi-isotropic coupling. The electrostatic interactions were modelled utilising the particle-mesh-Ewald algorithm [71] and a cutoff of 1.2 nm for the real-space contribution whereas the Lennard-Jones potentials were smoothly fixed to zero between 1 nm and 1.2 nm. Simulation were run with the GPU-accelerated Gromacs2020.1 [72].

#### The fitness function

The measurables extracted form the MD simulation are combined to one fitness function to measure the quality of the force field. The order parameter component of the fitness function (fit_OP_) reads

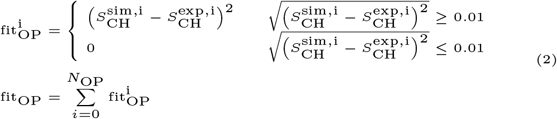

where the final fit_OP_ is obtained through a summation over *N*_OP_ individual order parameters. The fitness from the area per lipid (fit_A_) is

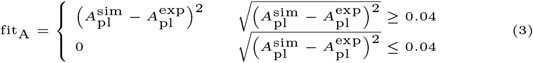

where 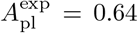and 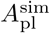 is the simulated area per lipid.

The final fitness is the weighted sum of the two target variable fitnesses fit=*m*_OP_*fit_OP_+*m*_A_*fit_A_. For the results presented here *m*_*OP*_ =10 and *m*_*A*_=15 were used. The partial definition of the component sets the fitness to zero when the simulated values are firmly within the uncertainty area of the experimental targets. For the *S*_CH_ values the experimental uncertainty is approximated as *±* 0.02 [38] whereas the target range for *A*_pl_ (0.60– 0.68 nm^2^) reflects the spread of values found in literature [47, 55, 56].

### Validation

Once a sufficient agreement with the experimental target observables was achieved, we performed a validation of the force field using a longer simulation (1-*µ*s) of the same 34-lipid system. First, we confirmed that optimization targets are reproduced within the experimental uncertainty in these longer simulations while accounting for simulated uncertainty obtained by first quantifying the order parameters of individual lipids over time, and then averaging over individual lipids, from which the standard error of mean is used as the error estimate. We then proceeded to quantify the NMR *R*_1_ rates and effective correlations times, as well as SAXS form factors from these longer simulations to validate the force field against observables not used as optimization targets. For this, the trajectories were preprocessed so that the bilayer resided at the center of the box in z-direction. SAXS form factors were calculated frame-vise from the electron density profiles (Eq. 12 in Ref. 45) and then averaged over the frames to obtain the final curves. The strategy detailed in Ref. 49 was used for the NMR observables quantifying the conformational dynamics.

For the best model, for which data are presented here, we also conducted a 1-*µ*s simulation of a larger (200 lipids) system to confirm the absence of finite size effects beyond those known to affect the simulated SAXS curves due to the membrane fluctuations [73].

### Self organized maps

Self organized maps [74] is a type of an artificial neural network which can be used to produce (typically) 2D visualization of large, multi-dimensional datasets. Each cell of the map contains a neuron (a vector) which has the dimensionality of the input data. During training, each input datapoint is assigned to the map cell whose neuron most closely resembles that input datapoint, and the neuron in that cell, as well as in the neurons in the neighbouring cells, are shifted towards the input. This will result in a map that preserves the input space topology.

For creating the self organized maps, we used lipid conformations from a 1-*µ*s simulation of the 34-lipid system. Each lipid was first translated so that their g_2_ carbon coincided with the coordinate zero and flipped so that the headgroup resided on the positive *z*-coordinate. Then the lipid was rotated in the *x*–*y* plane so that the *x*–*y* component of the g_2_–g_1_ bond coincided with the *x*-axis. These transformations preserve the lipid tilt with respect to the bilayer normal, which we view as a property of the lipid conformational state. The coordinates of the heavy atoms in the lipid where then used to create the self organized maps.

We divide POPC into 3 parts, 1) sn-1 chain, 2) sn-2 chain, and 3) headgroup+glycerol backbone, splitting the molecule from the g_2_ carbon so that this carbon does not belong to any of the three molecular fragments. We then proceed to train a self organized map of size 50 × 50 neurons for each of these fragments based on the heavy atom coordinates. In the training, the coordinates of 3400034 conformations are cycled though four times and the learning rate was exponentially decreased from 0.5 to zero. The maps have periodic boundary conditions.

When choosing the SOM parameters, we confirmed that further increasing the size of the map made no difference to the general features observed nor the rank plots [75]. Increasing the number of cycles through the data in training did not significantly decrease the quantization or topological errors. The self organized maps were created using Quicksom [76].

