## Supplementary material for "Autofocusing molecular simulations for integrative structural biology of broad conformational ensembles"

### Simulated vs experimental structures

#### *Lipid headgroup tilt*

Roberts and Redfield [1] have estimated from their $^{31}$P NMR experiments on binary POPC mixtures that the vector from the headgroup P to the $g_3$ hydrogens forms either a 41–44° or 67–72° angle with the bilayer normal. The two ranges come from the two solutions of the equation used to extract the angle from $^{31}$P NMR data. Our model indicates the average angle is 51°, and 37% of conformations are within 10° and 53% are within 15° of this average. Within range 41–44° are 5% and within 67–72° are 6% of the conformations. In the phosphatidylcholine headgroup model by Akutsu and Nakamori [2], proposed based on NMR and Raman spectroscopy data, the headgroup P-N vector has an angle of 18° with the membrane surface. This matches the average P-N orientation obtained from our model.

#### *Glycerol backbone orientation*

Figure S2 shows the orientations (with respect to the bilayer normal) for two bond vectors of the central glycerol carbon $C^{g_2}$. Interestingly, and unlike to what is observed in the existing crystal structures (see the thin vertical lines in Fig. S2A), the $C^{g_2}$–$O^{g_2}$ bond (in red) does not align in the membrane plane ($\theta = 90°$) as much as the $C^{g_2}$–$C^{g_1}$ bond (black) does. Thus we also chose to align our structures such that the projection of the $C^{g_2}$–$C^{g_1}$ (and not $C^{g_2}$–$O^{g_2}$) bond to the membrane ($xy$) plane is aligned with the $x$-axis.

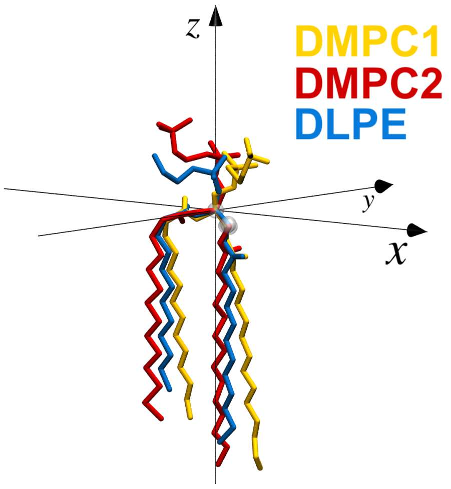


FIG. S1: X-ray crystal structures of DMPC [3] (orange and red) and DLPE [4] (blue) phospholipids. Centered on glycerol carbon $C^{g2}$ (origin), and rotated around bilayer normal ($z$) such that the projection of $C^{g2}$–$C^{g1}$ bond in the $xy$-plane aligns with the $x$-axis. In the visualization the $C^{g1}$ and $C^{g2}$ atoms are highlighted with white spheres.

Classic analysis of the PC headgroup conformation starts with the assumption (first presented by Seelig and Gally for PE headgroups [6]) that the $C^{g2}$–$C^{g3}$ bond preferably aligns with the bilayer normal, about which it wobbles due to thermal noise [7]. We find this assumption to hold fairly well: The most probable orientation of $C^{g2}$–$C^{g3}$ occurs at roughly 10° angle, with the 0° also well populated (Fig. S3A,C). The orientational distribution is markedly wide and, strikingly, symmetric with respect to the glycerol-backbone plane ($y = 0$) while being asymmetric with respect to the $x = 0$ plane, see Fig. S3A–C.

The average angle between the $C^{g2}$–$C^{g3}$ and bilayer normal is $31.1 \pm 0.2°$ (white circle in Fig. S3A,B), matching the $35 \pm 10°$ of the single characteristic conformation by Hong et al. [5]. It is worth noting that this average angle markedly differs from the angle (of roughly 10°) of the most probable orientation; the apparent contradiction is caused by the peculiarities of angular averaging, as demonstrated by Fig. S3D, which shows the ratio between the observed probability density and the probability density that would be observed in the case of a uniform (spherical)

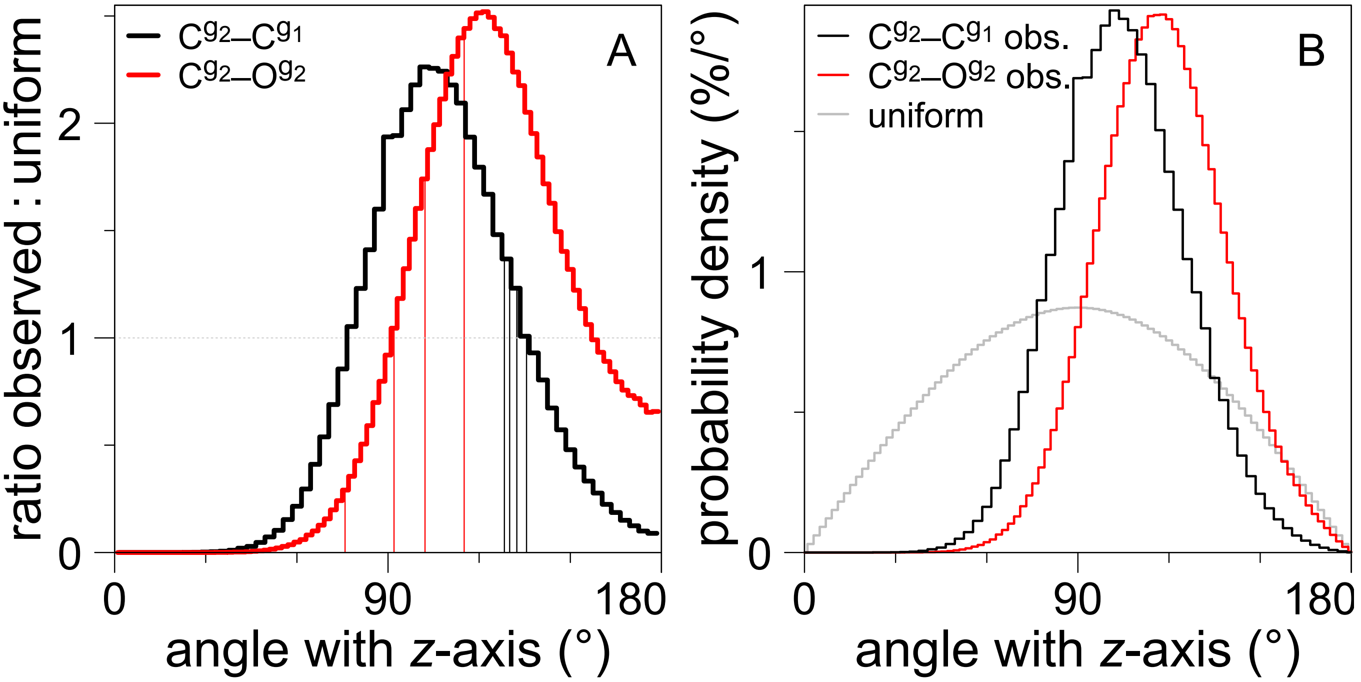


FIG. S2: Angle distributions between the bilayer normal (aligned with the $z$-axis) and two bond vectors: $C^{g_2}$–$C^{g_1}$ (in black) and $C^{g_2}$–$O^{g_2}$ (red). Panel (A) shows the ratio between the observed vector orientations and the case of no preferred orientation (uniform spherical distribution); panel (B) shows the observed distributions as well as how a uniform spherical distribution (in gray) would appear. The thin vertical lines in panel (A) indicate the angles reported in the literature for a DLPE crystal [4] (102° for $C^{g_2}$–$O^{g_2}$ / 128° for $C^{g_2}$–$C^{g_1}$), a DMPC crystal [3] (92° / 130° and 115° / 136°), and an NMR-consistent single conformation of DMPC in liquid-disordered state [5] (76° / 132°).

distribution.

To characterize the extent of angular excursions of the wobble, Seelig and Gally [6] introduced an order parameter for the $C^{g2}$–$C^{g3}$ bond, and estimated it to be 0.66; we find it to be $0.559 \pm 0.003$. For their estimate, they took an average order parameter of |0.22| for the $C^{g_3}$–H bonds (we find: $-0.237$), 109.5° for the $C^{g2}$–$C^{g3}$–H angle ($107.9 \pm 0.1°$, see Fig. S3E for the distribution), and assumed free rotation—or at least a threefold symmetric jump—to occur about $C^{g2}$–$C^{g3}$. The last assumption is the one that clearly disagrees with the autofocused simulations (Fig. S3F), resulting in the difference observed.

*Forking*

Occasionally, a peculiar feature where two C–H bond on same carbon have distinctly different order parameters (aka. forking), is detected. In POPC, this happens at the $g_1$ and sn-2 C2 carbons. Forking can originate from two scenarios: 1) the molecule has two long-lived (wrt. measuring time) conformations or distinct conformational subpopulations, each of the two separately having equivalent C–H bonds or 2) two C–H bonds with non-equivalent motions. Especially the molecular origin of the forking at sn-2 C2 gathered lot of debate [8, 9], with experimental evidence for both explanations. In particular, the existence of two distinct conformations in crystal structures favored option 1 [3]. However, scenario 2 finally emerged as winner when only one order parameter was observed after site-specific deuteriation of one of the C–H bonds [9]. Our model is in line with this and gives rather narrow, single-peak distributions for both C2 C–H angles (Fig. S4) instead of distributions shapes that would indicate two separate populations of angles for single C–H bond. Same is true for the $g_1$ hydrogen bonds. This exemplifies that caution should be exercised when using crystal structures as inspiration for interpretation of data in the liquid disordered phase.

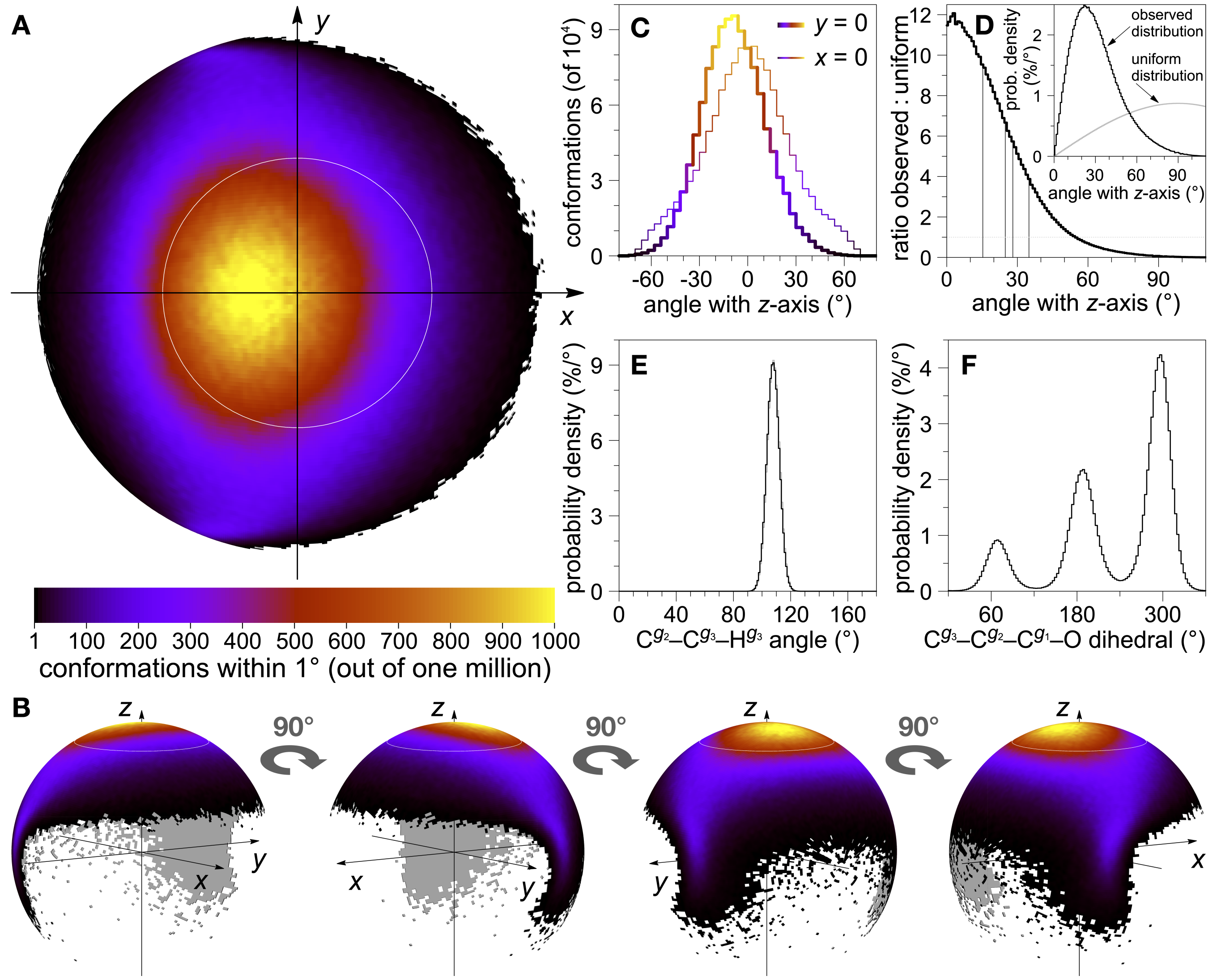


FIG. S3: The most probable orientation of the $C^{g2}$–$C^{g3}$ bond vector is almost parallel to the bilayer normal. (A) The observed distribution of $C^{g2}$–$C^{g3}$ vectors. Here $C^{g2}$ is in the origin, the bilayer normal ($z$-axis) points towards the viewer, and the $x$-axis is set to align with the $C^{g2}$–$C^{g1}$ vector's projection in the membrane ($xy$) plane. The white circle indicates the average angle ($31.1 \pm 0.2°$) between the $C^{g2}$–$C^{g3}$ bond vector and the bilayer normal. (B) Oblique views of the same spherical distribution. (C) Cuts of the spherical distribution with the $xz$-plane (thick line) and the $yz$-plane (thin); colouring as in panel A. (D) Ratio of two probability distributions for the angle between the $C^{g2}$–$C^{g3}$ bond vector and the bilayer normal: the one observed (black curve in inset) and the one expected if $C^{g2}$–$C^{g3}$ has a uniform spherical distribution (gray in inset). The thin vertical lines indicate the angles reported for a DLPE crystal [4] (15°), a DMPC crystal [3] (25° and 28°), see also Fig. S1, and an NMR-consistent single conformation of DMPC in liquid-disordered state [5] (35°). (E) Probability distributions of the $C^{g2}$–$C^{g3}$–$H^{g3}$ angles: one $H^{g3}$ in black, the other in gray. (F) Probability distribution of the $C^{g3}$–$C^{g2}$–$C^{g1}$–O dihedral angle.

#### *Tail conformations: leading chain and number of gauche conformations*

Pezeshkian *et al.* [10] considered angle of the O1–C1–C2–C3 dihedral ($\gamma_3$ dihedral for sn-1 and $\beta_3$ dihedral for sn-2 in Sundralingam notation, see Fig. S6) to be a marker of the leading chain: which ever chain had this dihedral mostly around 180° was the leading one. However, in our model both chains have similar probabilities (minimum) in the immediate vicinity of 180°. Sn-2 has a peak at 200° while sn-1 symmetrically switches between two maxima around the conformation at 180°, see Fig. S5 (top). This analysis alone is rather inconclusive. However, directly looking at the C1–C2 bond reveals that the sn-1 chain has high probability an angle around 180° with respect to the bilayer normal (simulation z-axis) and with respect to the $g_2$–$g_3$ (Fig. S5 bottom). The latter is analyzed to remove the effect of the lipid tilt. In contrast, the sn-2 chain has distributions more weighted towards 90°. Note, however, that

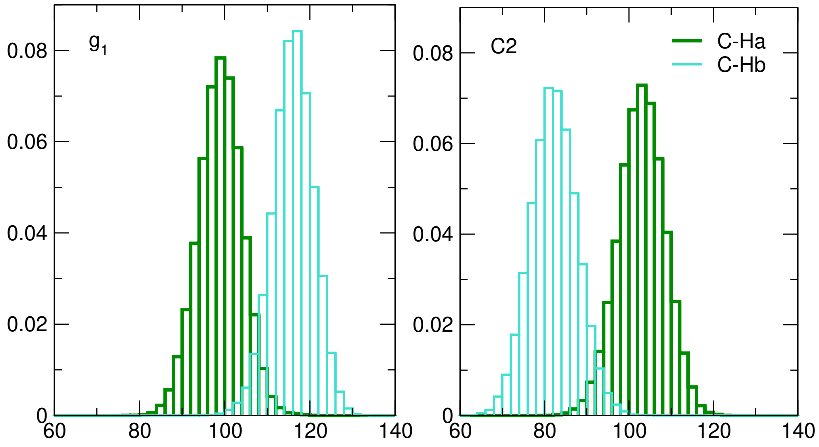


FIG. S4: Histograms of C-H bond angles $\theta$ with the z-axis for the carbons where forking occurs in POPC, $g_1$ (right panel) and oleoyl C2 carbon (left panel). Colors separate the two hydrogens attached to each carbon, denoted Ha and Hb.

both chains have a finite probability for conformations around 90° and 180°, indicating that sn-2 can occasionally adopt the leading chain conformation and sn-1 can exhibit conformations with C1–C2 bond parallel to the bilayer. Specifically, sn-1 chain C1–C2 has a peculiar bimodal angle distribution of with respect to $g_2$–$g_3$.

In our analysis of tail conformations, we use the following angle limits to define gauche and trans dihedral conformations: 0-120° for gauche+, 120-240° for trans, and 240-360° for gauche-. The experimental estimates for number of simultaneous gauche conformations in saturated acyl chains have been usually above 3: Ref. [11] gives 3-6 simultaneous gauche bonds in L-$\alpha$-dipalmitoyllecithin (using NMR), whereas Ref. [12] estimates 3.6-4.2 and Ref. [13] 3.7±0.2, both in DPPC using FTIR. The saturated palmitoyl tail in our model has on average 4.02 simultaneuous gauche conformations which is in line with the experiments. More specifically, Mendelson *et al.* [12] measured the DPPC dihedrals around carbon 4 to be in gauche conformation 20.7± 4.2% of time, carbon 6 32.3±2.3% and carbon 10 19.7±0.8% of time. Our POPC model gives 14% gauche for 3'–4' dihedral, 29% for 4'–5', 30% for 5'–6', 32% for 6'–7' and 32% for both 9'–10' and 10'–11' dihedrals. Only the value for the last carbon is clearly larger than the experimental one. While we would expect the conformations of the oleoyl tail in POPC to be rather independent of the presence of the double bond at the palmitoyl tail, introduction of such double bond is known to increase disorder in the membrane and consequently alter membrane properties by, for example, causing an increase in the area per lipid [14, 15]. By creating more space, this would promote the number of gauche conformations in the saturated tail of POPC compared to those of DPPC [16] (at the same temperature), for which reason the comparison between the two should be mostly qualitative.

Similarly, there is experimental evidence that having a double bond in a lipid tail may increase number of gauche conformations around that double bond compared to a saturated chain [16]. This is in line with our model, where percentage of gauche conformations from 4'-5' bond onwards is ~30% in the saturated tail but around the double bond peaks to 40%. However, due to the underlying complicating assumptions in the experiments [16], we refrain from further comparison. The main experimental indication of such double bond remains the dip in the absolute value of the order parameters, which is a feature usually well reproduced in MD simulations [15, 17, 18], and which signals the disorder-inducing impact the double bond has on the membrane [15].

#### *Comparison to consensus conformations from NMR and scattering experiments*

Here we present the simulated conformations in terms of dihedral angles along with the comparison to the consensus structures, extracted from scattering experiments and predicted by the early pioneers of lipid NMR, see Table S1. These structures cover mostly the headgroup, extending at most to the parts of the lipid tails attached to the glycerol backbone. The used dihedral notation follows the Sundralingam convention summarized in Fig. S6. The full dihedral distributions for the entire molecule are given in Fig. S7. In general, the fractions of conformations in the simulation which fully agree with the structures predicted based on the experimental data remain low. In fact, no conformations were present in the simulations for which all the dihedrals were in agreement with the X-ray structures: Despite the high agreement of some individual dihedrals with the experimental consensus, it is highly improbable that all

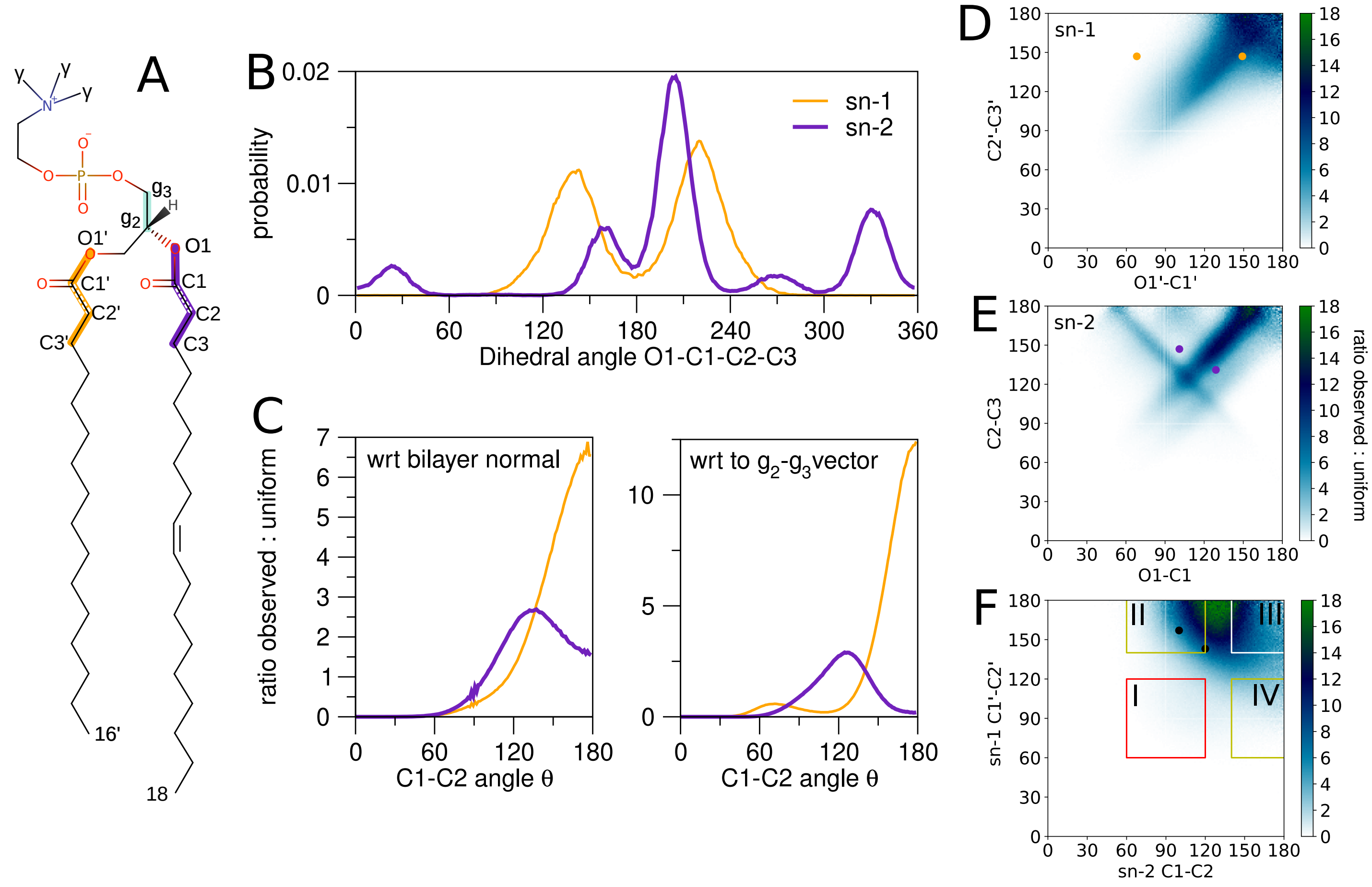


FIG. S5: Insight into the leading chain. A) The vectors and the dihedral angles along with the carbon labelling. B) O1-C1-C2-C3 dihedral for both lipid tails (top panel). Both tails have dihedrals with nonzero probabilities around 180°. C) The probability distribution for angle between bilayer normal and C1-C2 vector (left) and between $g_2$–$g_3$ and C1-C2 vectors (right). Distributions are more weighted towards 90° for sn-2 chain and towards 180° for sn-1. D-F) 2D probability maps for D) sn-1 C2–C3 vs O1–C1 angles, E) sn-2 C2–C3 vs O1–C1 angles with respect to the z-axist and F) sn-1 C1–2 vs. sn-2 C1–C2 angles with respect to the z-axis. The colored dots denote the sn-1 (yellow) and sn-2 (purple) angles in the crystal conformations taken from Ref. [3]. Bottom panel gives the 2D probability distribution for sn-1 C1–C2 vs sn-2 C1–C2 angles with respect to the z-axis. The black dots denote the angles in two crystal conformations taken from Ref. [3], see also Fig. S1. Four areas are marked in the plots. I: both chains have a kink conformation (angle within the window 90° ± 30°), II: sn-1 leading (170° ± 30°), sn-2 kink. III: both chains leading. IV: sn-1 kink, sn-2 leading.

angles match the experimental prediction simultaneously. For individual dihedrals, the highest agreement with the experimentally predicted angle and that observed in the simulation is just under the fraction of 0.7 of conformations.

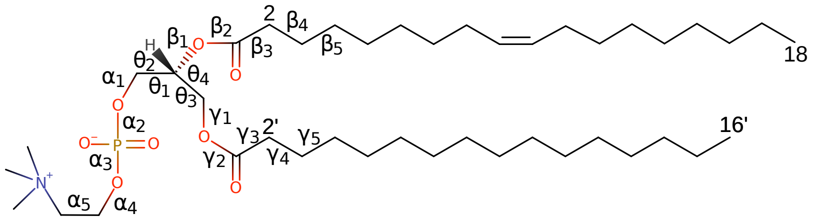

FIG. S6: Sundaralingam notation for dihedrals involving N, C and ester oxygens.

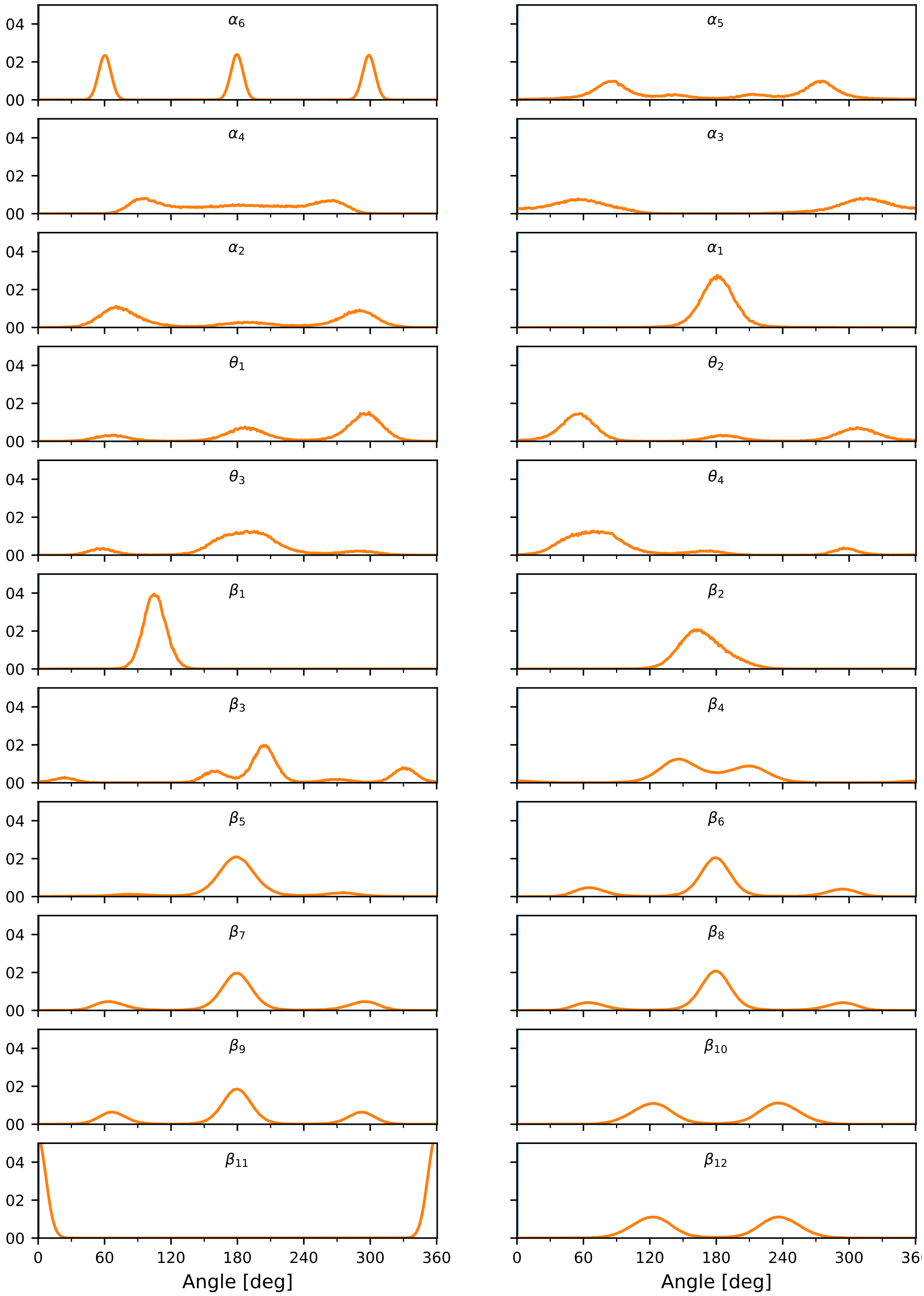

$\alpha_6$
$\alpha_5$
$\alpha_4$
$\alpha_3$
$\alpha_2$
$\alpha_1$
$\theta_1$
$\theta_2$
$\theta_3$
$\theta_4$
$\beta_1$
$\beta_2$
$\beta_3$
$\beta_4$
$\beta_5$
$\beta_6$
$\beta_7$
$\beta_8$
$\beta_9$
$\beta_{10}$
$\beta_{11}$
$\beta_{12}$
04
02
00
0
60
120
180
240
300
360
Angle [deg]

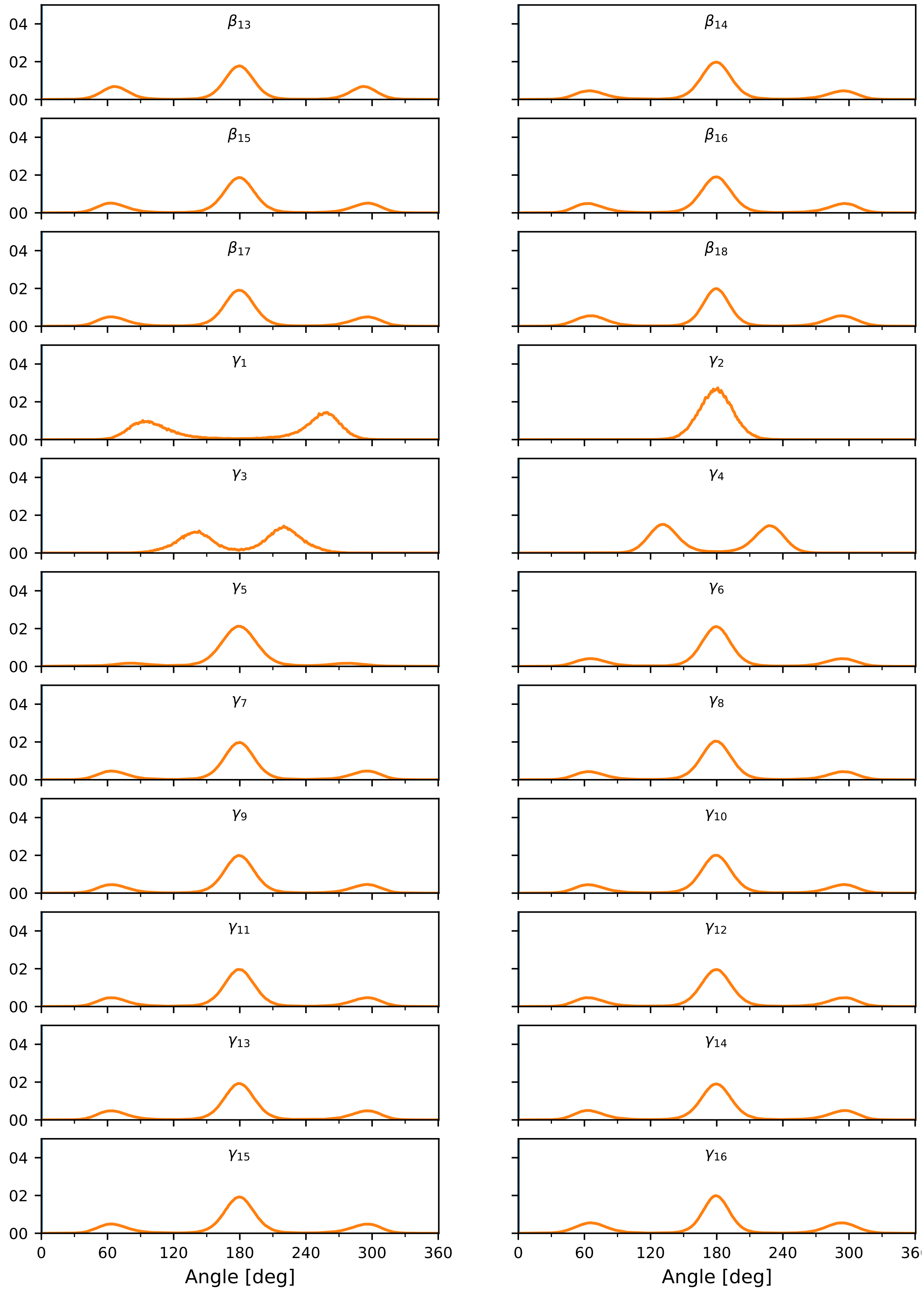


FIG. S7: Dihedral angle distributions of the calibrated model 598-3. The labelling of dihedrals follows the Sundaralingam notation presented in Fig. S6.

| Experiment (lipid)/Ref | | $\alpha_1$ | $\alpha_2$ | $\alpha_3$ | $\alpha_4$ | $\alpha_5$ | $\theta_1$ | $\theta_2$ | $\theta_3$ | $\theta_4$ | $\beta_1$ | $\beta_2$ | $\beta_3$ | $\beta_4$ | $\gamma_1$ | $\gamma_2$ | $\gamma_3$ | $\gamma_4$ | MD fraction |
|---|---|---|---|---|---|---|---|---|---|---|---|---|---|---|---|---|---|---|---|
| NMR (DPPC)/[7] | angle | 170 | 300 | 296 | 215 | 81 | - | - | - | - | - | - | - | - | - | - | - | - | 0.002 |
| | MD | 0.569 | 0.214 | 0.183 | 0.115 | 0.222 | - | - | - | - | - | - | - | - | - | - | - | - | |
| NMR (DPPC)/[7] | angle | 190 | 60 | 64 | 145 | 279 | - | - | - | - | - | - | - | - | - | - | - | - | 0.002 |
| | MD | 0.570 | 0.213 | 0.182 | 0.117 | 0.216 | - | - | - | - | - | - | - | - | - | - | - | - | |
| NMR (DPPC)/[2] | angle | - | - | 301 | 210 | 83.2 | - | - | - | - | - | - | - | - | - | - | - | - | 0.008 |
| | MD | - | - | 0.197 | 0.115 | 0.226 | - | - | - | - | - | - | - | - | - | - | - | - | |
| NMR (DPPC)/[2] | angle | - | - | 51 | 120 | 83.2 | - | - | - | - | - | - | - | - | - | - | - | - | 0.018 |
| | MD | - | - | 0.208 | 0.136 | 0.226 | - | - | - | - | - | - | - | - | - | - | - | - | |
| NMR (DPPC)/[2] | angle | - | - | 56 | 130 | 89.5 | - | - | - | - | - | - | - | - | - | - | - | - | 0.011 |
| | MD | - | - | 0.203 | 0.121 | 0.218 | - | - | - | - | - | - | - | - | - | - | - | - | |
| NMR (DPPC)/[2] | angle | - | - | 303 | 130 | 82.7 | - | - | - | - | - | - | - | - | - | - | - | - | 0.008 |
| | MD | - | - | 0.202 | 0.121 | 0.225 | - | - | - | - | - | - | - | - | - | - | - | - | |
| NMR (DPPC)/[2] | angle | - | - | 313 | 120 | 73.7 | - | - | - | - | - | - | - | - | - | - | - | - | 0.008 |
| | MD | - | - | 0.207 | 0.136 | 0.192 | - | - | - | - | - | - | - | - | - | - | - | - | |
| NMR (DPPC)/[2] | angle | - | - | 319 | 120 | 69.3 | - | - | - | - | - | - | - | - | - | - | - | - | 0.006 |
| | MD | - | - | 0.196 | 0.136 | 0.163 | - | - | - | - | - | - | - | - | - | - | - | - | |
| NMR (POPC)/[19] | angle | - | - | 62 | 194 | 308 | - | - | - | - | - | - | - | - | - | - | - | - | 0.002 |
| | MD | - | - | 0.189 | 0.120 | 0.065 | - | - | - | - | - | - | - | - | - | - | - | - | |
| NMR (POPC)/[19] | angle | - | - | 299 | 194 | 52 | - | - | - | - | - | - | - | - | - | - | - | - | 0.003 |
| | MD | - | - | 0.192 | 0.120 | 0.066 | - | - | - | - | - | - | - | - | - | - | - | - | |
| X-ray (DMPC)/[20] | angle | 163 | 62 | 68 | 143 | 296 | 58 | 177 | 182 | 63 | 82 | 172 | 279 | 45 | 183 | 168 | 187 | 178 | 0 |
| | MD | 0.416 | 0.226 | 0.167 | 0.117 | 0.123 | 0.205 | 0.068 | 0.336 | 0.215 | 0.094 | 0.409 | 0.020 | 0.002 | 0.017 | 0.540 | 0.079 | 0.027 | |
| X-ray (DMPC)/[20] | angle | 177 | 286 | 313 | 210 | 54 | 168 | 280 | 166 | 51 | 120 | 179 | 226 | 67 | 102 | 176 | 180 | 180 | 0 |
| | MD | 0.661 | 0.252 | 0.207 | 0.115 | 0.073 | 0.151 | 0.104 | 0.307 | 0.182 | 0.264 | 0.418 | 0.083 | 0.001 | 0.298 | 0.661 | 0.066 | 0.026 | |
| X-ray (DMPC)/[3] | angle | 162 | 68 | 63 | 139 | 309 | 58 | - | 184 | - | 76 | 180 | 273 | 63 | 187 | - | - | - | 0 |
| | MD | 0.393 | 0.252 | 0.186 | 0.117 | 0.061 | 0.205 | - | 0.334 | - | 0.034 | 0.419 | 0.023 | 0.001 | 0.018 | - | - | - | |
| X-ray (DMPC)/[3] | angle | 170 | 284 | 314 | 199 | 64 | 169 | - | 175 | - | 124 | 169 | 234 | 63 | 103 | - | - | - | 0 |
| | MD | 0.569 | 0.247 | 0.206 | 0.117 | 0.127 | 0.155 | - | 0.334 | - | 0.187 | 0.401 | 0.028 | 0.001 | 0.296 | - | - | - | |
| NMR (DMPC)/[5] | angle | 110 | 170 | - | - | - | 185 | - | 265 | - | 145 | 110 | - | - | 290 | 180 | - | - | 0 |
| | MD | 0.004 | 0.061 | - | - | - | 0.172 | - | 0.021 | - | 0.019 | 0.007 | - | - | 0.070 | 0.677 | - | - | |

TABLE S1: Frequency of experimentally predicted conformations found in the calibrated MD simulation. First column gives the experimental system and a reference. Each row marked "angle" defines a conformation (aka. a set of dihedral angles, columns) suggested by the experiment in degrees. The row underneath (denoted "MD") gives the fraction of that individual dihedral angle found in the MD simulation using calibrated model. Last column gives the total fraction of the experimental conformation (each defined dihedral occurring simultaneously) in simulations. Fractions calculated based on $\pm 15^\circ$ tolerance around given angle value. Hyphen denotes no information for angle. The used Sundaralingam notation [21] for dihedral angles is elaborated in Fig. S6.

*Dihedral isomerization rates*

In Figure S8 we visualize the angle of the tail dihedrals as a function of time for a selected lipid in a bilayer. The visualization is done for a snipped of 25 ns from the simulation with a sampling rate of 10 ps.

*Insights from the self organized maps*

Figure S9 shows paths drawn on the SOMs depicting the conformational space of the lipid headgroup and the two tails separately. As in the analogous figure in the main text (Fig. 3), the paths drawn via the peaks and valleys of the SOMs do not exhibit a large variation in energy while still having fairly large conformational variation, evident from the visualization of the SOM-cell average conformations and conformations best matching the SOM cell from the trajectory along the drawn path. That is, the conformational space does not contain distinct states separated by high barriers but is rather continuous with the observed conformations all within energy variation of $\sim 3k_{\mathrm{B}}T$. This further evidences how the smoothness of the energy landscape and the lack of distinct conformational states makes the describing the conformational space using one or few example conformations misleading.

Figure S10 shows the U-matrices for the molecular SOMs and the clustering of conformations done based on these U-matrices on top of the SOM: areas of low U-matrix value (similar conformations between neighbouring cells) were grouped together whereas cluster boundaries follow areas of higher U-matrix values (higher conformational distance between cells). The right column shows the conditional probability matrices where created based on this clustering. These show very little conformational correlation between the clusters: the molecular fragments are conformationally fairly independent of each other. If one chooses a conformational cluster for a fragment presented in the x-axis, one always encounters a similar probability distribution of conformations occurring for the fragment in the y-axis.

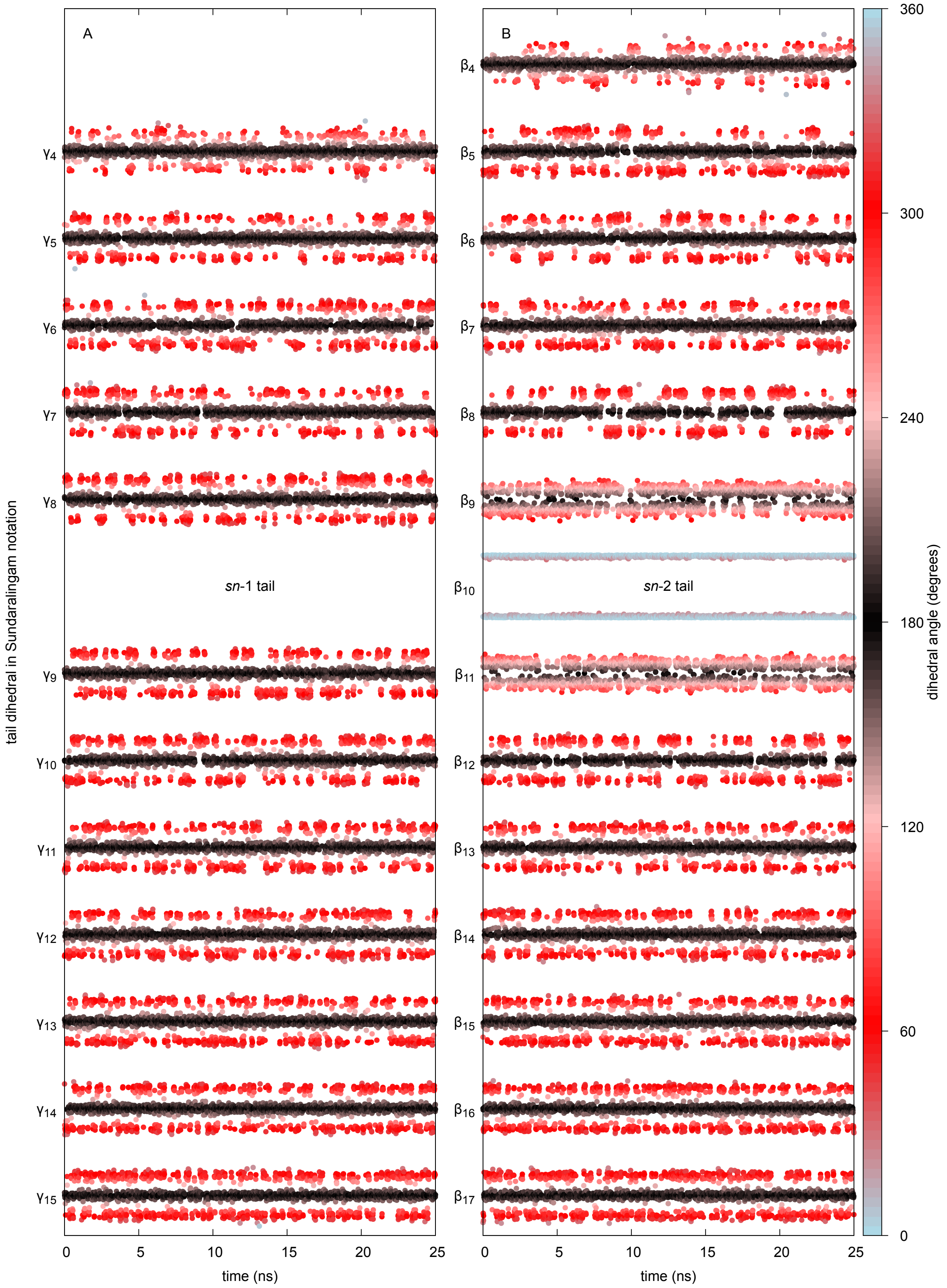


FIG. S8: Tail dihedral angle isomerization dynamics of a single POPC during a 25-ns simulation windown. (A) The palmitoyl (*sn*-1) tail, (B) the oleoyl (*sn*-2) tail; the labelling of dihedrals follows the Sundaralingam notation (Fig. S6). The point color indicates the dihedral angle value at the given time (sampling rate 10 ps); consequently, a trans state appears as black and gauche states red.

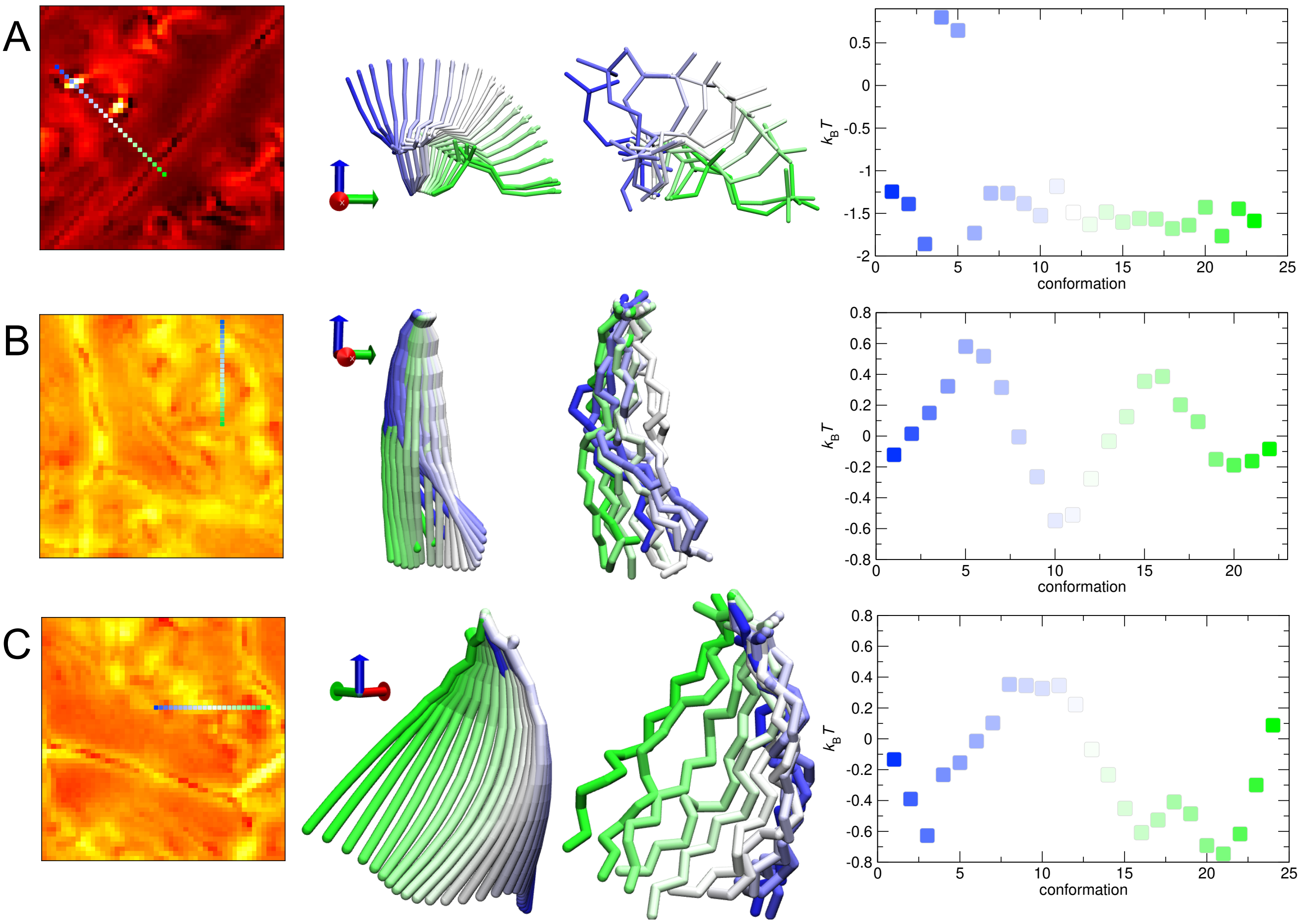


FIG. S9: Visualizing conformations of (A) head group, (B) sn-2 tail, and (C) sn-1 tail using self organized maps (SOM). Left panels: paths drawn on top of self organised maps that show the conformational free-energy landscape. Middle panels: the conformations corresponding to the drawn paths for the SOM-cell average conformation (middle left) and the conformation from the trajectory that is closest to the SOM-cell average conformation (middle right). The $z$-axis (blue) aligns with the bilayer normal, the $x$-axis (red) with the $C^{g2}$–$C^{g1}$ vector's projection in the membrane ($xy$) plane. Right panels: free energies in units of $k_BT$ of the SOM-cells along the drawn path.

Conversely, for each conformation of the molecular fragment on the y-axis, there is no clearly favored cluster for the fragment in the x-axis.

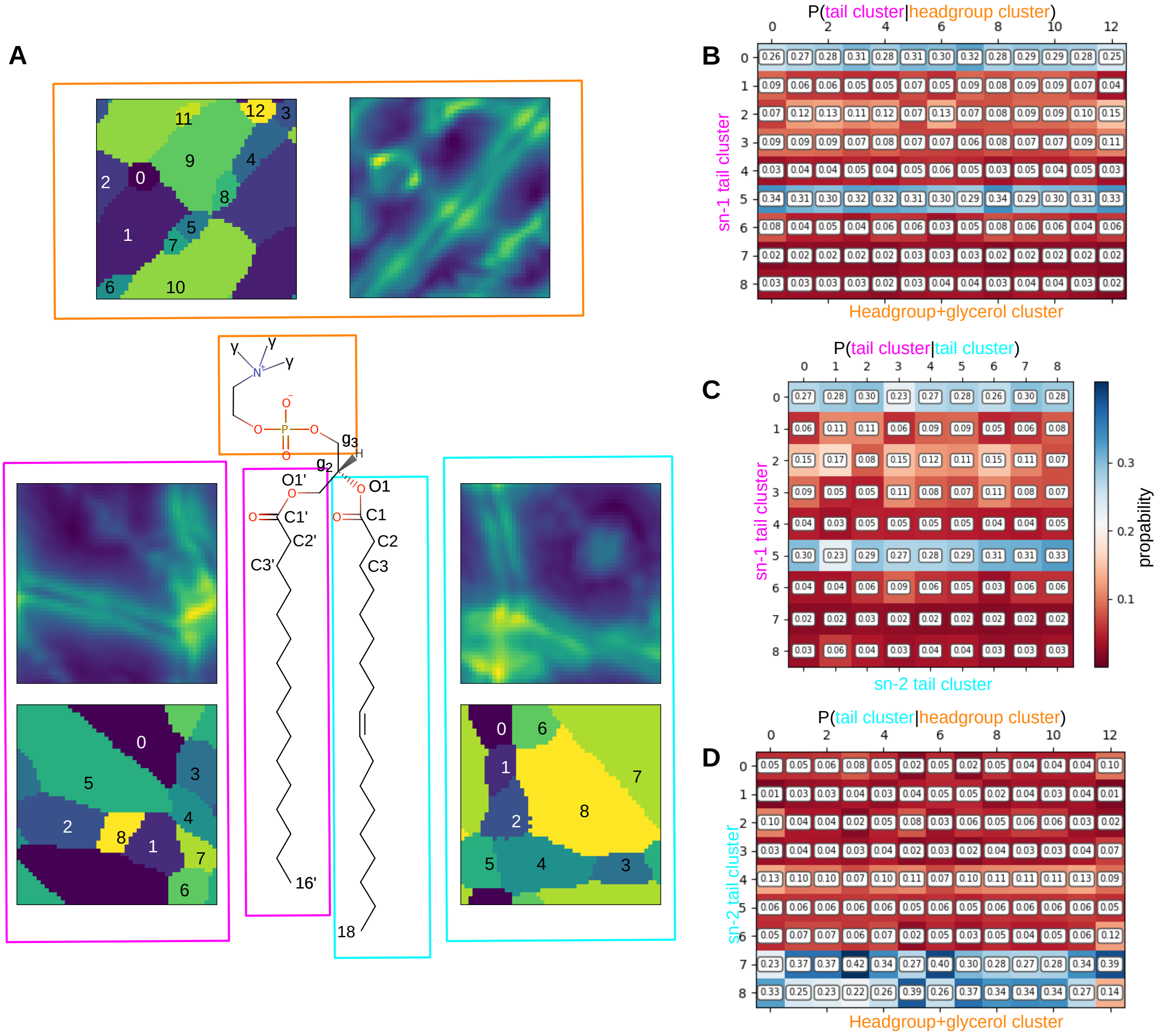


FIG. S10: A) the U-matrices of the self organized maps for different molecular fragments (colored boxes) next to the numbered conformation clusters defined based on the U-matrix. The U-matrices depict the average distance to neighboring SOM cells, the brighter yellow colors denoting cells where conformations differ more from those assigned to the neighbouring cells (high average distance). B-D) the conditional probability matrices for conformational clusters for different combinations of molecular fragments, color coding according to panel A). B) sn-1 tail clusters vs headgroup clusters, C) sn-1 tail clusters vs sn-2 tail clusters, D) sn-2 tail clusters vs headgroup clusters. Note that the matrix columns add up to one.

[20] Marsh, D. *Handbook of Lipid Bilayers* (CRC Press, 2013), 2nd edn.

[21] Sundaralingam, M. Discussion paper: molecular structures and conformations of the phospholipids and sphingomyelins. *Annals of the New York Academy of Sciences* **195**, 324–355 (1972).
